# MLL1 Methyltransferase Activity is Regulated by Distinct Nucleosome Binding Modes

**DOI:** 10.1101/2021.04.13.439652

**Authors:** Alex Ayoub, Sang Ho Park, Young-Tae Lee, Uhn-Soo Cho, Yali Dou

**Author notes:** Equal contribution (Alex Ayoub, Sang Ho Park). Co-correspondence, Lead contact.

## Abstract

Here we solve the single particle cryoEM structure for the MLL1 complex with nucleosome core particle (NCP) carrying histone H3 lysine 4 to methionine mutation. The MLL1 complex displays significant rotational dynamics on the NCP, a feature distinct from the yeast SET1 complex. We identified two major binding modes of the MLL1 complex on the NCP. Both binding modes anchor on the NCP through ASH2L, but they differ drastically with regard to where the MLL1 SET domain and RbBP5 bind. We show that one of the binding modes is catalytically inactive since disrupting interactions unique to this binding mode does not affect overall MLL1 activity in an NCP-specific manner. Interestingly, the inactive binding mode is in a configuration similar to that of the ySET1- NCP complex, which is intrinsically inactive on an unmodified NCP. The high rotational dynamics of the MLL1 complex as well as distinction between MLL and yeast SET1 complexes may reflect the necessity for loci-specific regulation of H3K4 methylation states in higher eukaryotes.

## INTRODUCTION

The Mixed Lineage Leukemia 1 (MLL1/KMT2A) belongs to the MLL/SET1 family of histone H3 lysine 4 (H3K4) methyltransferases. It is able to deposit H3K4 methylation (H3K4me), which demarcates active transcription and recruits basal transcriptional machinery (1-4). Heterozygous mutations for the MLL/KMT2 family enzymes are found in multiple human congenital diseases (5). MLL/KMT2s are also among the most frequently mutated genes in cancer (5). Through a highly conserved C-terminal catalytic SET domain, MLL1 interacts with an essential core complex of proteins, including RbBP5, WDR5, ASH2L, and DPY30 (MLL1^RWSAD^, MLL1.com). This conserved core complex is essential for efficient catalysis of mono-, di-, and tri-methylation of H3K4 (6-9). Recently, two groups, including ours, reported the cryoEM structures of the MLL1 core complex bound to the nucleosome core particles (NCPs) (10,11), illustrating how the MLL1 complex may engage its physiological substrate on chromatin. However, the two MLL1-NCP structures vary significantly in terms of how the MLL1 complex orients on the NCP. In Xue et al. (11), the MLL1 SET domain resides near the α2 and α3 helices of H2A with RbBP5 and ASH2L binding at NCP superhelical loop (SHL) 2.5 and 7, respectively (Figure S1A) (11). This orientation on the NCP is similar to that of the yeast SET1 complex (Figure S1C) (12,13), which is in a constitutively repressed state in the absence of histone H2B K123 ubiquitylation (H2BK120 ubiquitylation, or H2BK120ub, in human) (14,15). In Park et al. (10), we show that the MLL1 complex adopts a conformation rotating ∼26° counterclockwise with ASH2L bound at SHL 7 as a hinge, and RbBP5 reorienting to SHL 1.5 (Figure S1B). This conformation was also captured as a minor population in Xue et al. (11). In the second conformation, the MLL1 SET domain sits close to the nucleosome dyad, with almost symmetric access to both histone H3 tails for optimal processivity (Figure S1B) (10). Distinct binding modes of the MLL1 complex on the NCP raise the question of whether both MLL1-NCP configurations are functionally active on the NCP.

## RESULTS

To examine the functional significance of different MLL1-NCP interaction modes, we further stabilized the MLL1-NCP complex by using the NCP containing histone H3 lysine 4 to methionine (H3K4M) mutation (NCP^K4M^) (Figure S2A). Previous studies show that K to M mutation often traps the SET domain in a bound state (16) and the H3K4M mutant is able to stably interact with the SET domain (17). Indeed, the gel shift assay shows that the MLL1 complex bound more tightly to the NCP^K4M^ as compared to the wild-type NCP (Figure S2B). Next, we resolved the single particle cryoEM structure of the MLL1 complex with the NCP^K4M^. We obtained a total of 808,836 particles of the MLL1-NCP^K4M^ complex from 2,377 micrographic images. It captured two major populations of the MLL1-NCP^K4M^ complex from 3D classification (Figure 1A and S3). The structure of these two populations, i.e., mode 1 (29.2 %) and mode 2 (48.5 %), were determined at 4.76 Å and 4.02 Å resolution, respectively (Figure S4, S5 and Table 1). In mode 1 (MLL1-NCP^mode 1^), the MLL1 complex binds diagonally across the nucleosome disc, with interactions driven by RbBP5/NCP^SHL2.5^ and MLL1^SET^/histones^H3-H2A^ contacts (Figure 1A and S3A). In mode 2 (MLL1- NCP^mode 2^), the MLL1 complex binds at the edge of the NCP via RbBP5/NCP^SHL1.5^ and Ash2L/NCP^SHL7^ anchors (Figure 1B and S3B). These two modes of MLL1-NCP^K4M^ structures overlay nicely with previously reported MLL1-NCP structures (Figure 1C) (10,11). The MLL1- NCP^K4M^ mode 1 structure is similar to that reported by Xue et al. (11) as well as the ySET1-NCP structure (Figure 1B) (12,13) while the MLL1-NCP^K4M^ mode 2 structure is similar to the one reported by Park et al. (10). These results show that the MLL1 complex indeed displays significant rotational dynamics on the NCP, which is distinct from the yeast SET1-NCP complex (12,18).

**Figure 1.**
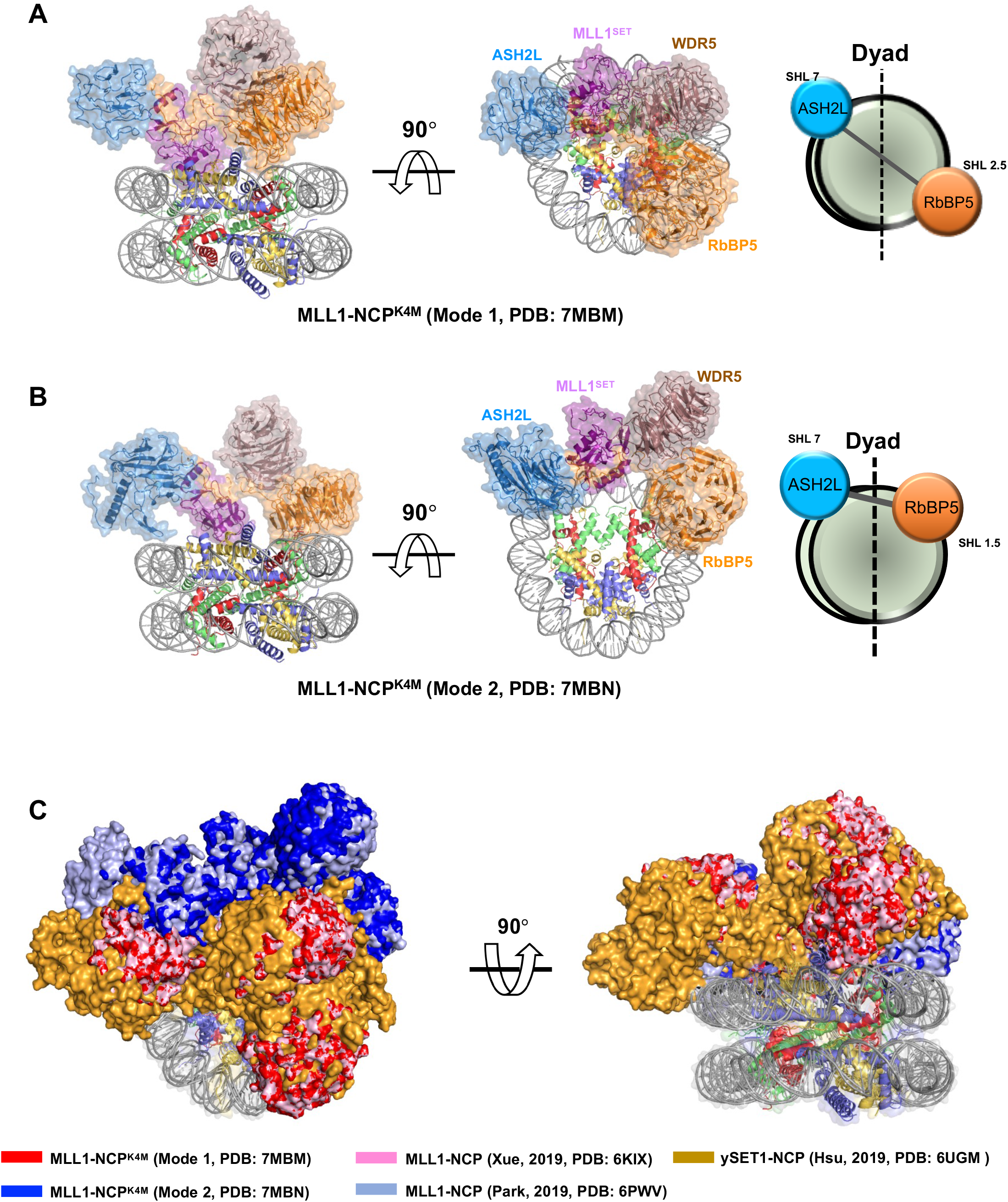
The Cryo-EM structures of the MLL1-NCP^K4M^. **a**, Front and top view of the MLL1- NCP^K4M, mode 1^ structure (PDB: 7MBM, EMDB: 23738). Right, cartoon model to show orientation of the MLL1 complex on the NCP. ASH2L (blue) and RbBP5 (orange) anchor the complex at SHL7 and SHL2.5 of the NCP, respectively. **b**, Front and top view of the MLL1-NCP^K4M, mode 2^ structure (PDB: 7MBN, EMDB: 23739). Right, cartoon model to show orientation of the MLL1 complex on the NCP. ASH2L and RbBP5 anchor the complex at SHL7 and SHL1.5 of the NCP, respectively. In both a and b, dyad axis is shown dashed line. **c**, Top (left) and front (right) views of aligned cryoEM structures of MLL1^RWSAD^-NCP from Xue et al. (pink, PDB: 6KIX, EMDB: EMD-0694), MLL1^RWSAD^-NCP from Park et al. (pale blue, PDB: 6PWV, EMDB: EMD-20512), ySET1-NCP from Hsu et al., 2019 (PDB: 6UGM, EMDB: EMD- 20765, goldenrod), MLL1-NCP^K4M, mode 1^ (red) and MLL1-NCP^K4M, mode 2^ (blue).

**Table 1.** CryoEM Data Collection, Refinement, and Validation Statistics

|  | MLL1<sup>RWSAD</sup>-NCP<sup>K4M</sup>,<br>Mode 1<br>(EMD-23738)<br>(PDB: 7MBM) | MLL1<sup>RWSAD</sup>-NCP<sup>K4M</sup>,<br>Mode 2<br>(EMD-23739)<br>(PDB: 7MBN) |
| --- | --- | --- |
| **Data Collection and Processing** |  |  |
| Magnification | 130,000 |  |
| Voltage (kV) | 300 |  |
| Electron exposure (e-/Å<sup>2</sup>) | 53.4 |  |
| Defocus range (μm) | -1.0 to -2.5 |  |
| Pixel size (Å) | 1.06 |  |
| Symmetry imposed | C1 |  |
| Initial particle images (no.) | 808,836 |  |
| Final particle images (no.) | 30,322 | 30,847 |
| Map resolution (Å) | 4.76 | 4.02 |
| FSC threshold | 0.143 | 0.143 |
| **Refinement** |  |  |
| Initial model used (PDB code) | 6KIX | 6PWV |
| Model resolution (Å) | 6.8 | 6.6 |
| FSC threshold | 0.5 | 0.5 |
| Map sharpening *B* factor (Å<sup>2</sup>) | -135.69 | -88.62 |
| **Model composition** |  |  |
| Non-hydrogen atoms | 19,757 | 20,222 |
| Protein residues | 1,747 | 1,806 |
| Nucleotides | 290 | 292 |
| Ligands | - | - |
| ***B* factor (Å<sup>2</sup>)** |  |  |
| Protein | 234.84 | 31.50 |
| Nucleotide | 109.84 | 9.31 |
| Ligand | - | - |
| **Rmsds** |  |  |
| Bond lengths (Å) | 0.006 | 0.005 |
| Bond angles (°) | 0.835 | 0.669 |
| **Validation** |  |  |
| MolProbity score | 2.01 | 2.60 |
| Clashscore | 9.39 | 37.16 |
| Poor rotamers (%) | 0 | 1.50 |
| **Ramachandran plot** |  |  |
| Favored (%) | 91.26 | 93.81 |
| Allowed (%) | 8.74 | 6.19 |
| Disallowed (%) | 0 | 0 |

Closer examination of two distinct MLL1-NCP^K4M^ binding modes reveals two major differences. In MLL1-NCP^mode 2^, we identified an arginine quartet (R220, R251, R272, and R294; Quad-R), an A-loop (_236_DGEPE_240_), and an I-loop (_193_TGTSNT_198_) in RbBP5 that are important for the NCP interaction (Figure 2A). They make close contact with the phosphate backbone of DNA as well as core histone H3 and H4, consistent with our previous report (10). In MLL1-NCP^mode 1^, while ASH2L retains binding near SHL 7, RbBP5 rotates clockwise to SHL 2.5 (Figure 2B). This rotation partially reorients the Quad-R motif and I-loop, thereby breaking most of their interactions shown in mode 2 (Figure 2B). Both R272 and R294 of Quad-R are detached from the DNA backbone interactions. Instead, a small loop (_294_RGE_296_) between WD40 domains 5 & 6 of RbBP5, specifically R294, makes a productive charged-charged interaction with E74 of H4 α2 helix (Figure 2B, loop 2). The essential I-loop is completely displaced from the core histones and instead, resides above DNA at SHL 2.5. Since the I-loop lacks positively charged residues, it is unlikely to interact with the DNA phosphate backbone. Instead of Quad-R and A/I-loops interactions, RbBP5 in MLL1-NCP^mode 1^ mainly interacts with α3 and αC helices of H2B through a highly conserved amphipathic loop, _248_LVNR_251_ (Figure 2B, Loop 1). The second major difference in two binding modes of the MLL1-NCP^K4M^ complex is the position of MLL1^SET^. In MLL1-NCP^mode 1^, the MLL1 complex rotates clockwise, compared to MLL1-NCP^mode 2^, leading to extensive interactions between MLL1^SET^ and the nucleosome disc. An extended helical patch (3806-21, including M3812, L3814, and M3818) in MLL1^SET^ make hydrophobic interactions with the α2 (N73) and α3 (L85) helices and C-terminus (L108 and P109) of H2A (Figure 2C). There is also an electrostatic interaction between an arginine anchor (R3821) of SET-N and D72 of the α2 helix of H2A (Figure 2C), a common feature in many protein-NCP complexes (19-23). These interactions are specific for MLL1-NCP^mode 1^.

**Figure 2.**
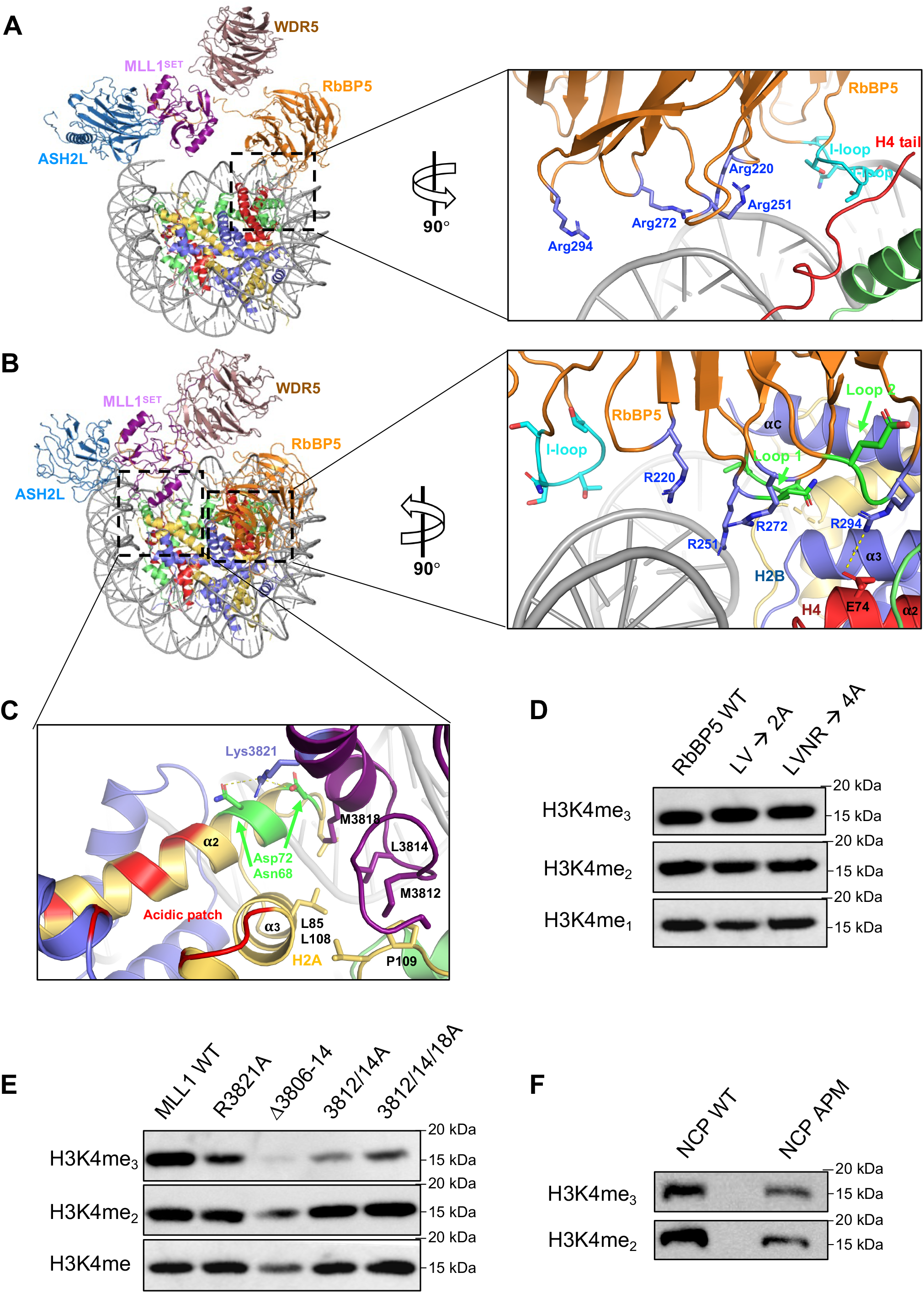
Unique interaction motifs in the MLL1-NCP^K4M, mode 1^ structure are dispensable for NCP-specific activities. **a**, Left, top view of the MLL1-NCP^K4M, mode 2^ structure. Right, inset shows Quad-R (shown in blue) and I-loop (cyan) that are engaged in the NCP interactions. **b**, Left, top view of the MLL1-NCP^K4M, mode 1^ structure. Right, inset shows that Quad-R (blue) and I-loop (cyan) dis-engage from the NCP in this configuration. New interactions involving RbBP5 _248_LVNR_251_ (Loop 2, green) are highlighted. **c**, Inset from **b** that shows the hydrophobic interface between MLL1 SET-N 3806-3821 (purple) and the α2, α3 and C-terminal helices (yellow) of H2A. Salt bright and polar contacts between R3821 (blue) of MLL1 and residues from H2A are shown. **a-c**, colors are as follows: ASH2L (light blue), MLL^SET^ (purple), WDR5 (tan) and RbBP5 (orange). **d-f**, *In vitro* histone methyltransferase assay using proteins indicated on top. All assays were carried out using the NCP as the substrate. **d**, RbBP5 and RbBP5 mutants; **e**, MLL1 or MLL1 mutants; **f**, NCP^WT^ or acidic patch mutant (NCP^APM^), were used in the reaction. Antibodies used in the immunoblot were indicated on left.

We previously showed that the MLL1-NCP^mode 2^ is functionally active and mutation of key residues, in the I-loop, A-loop, or arginine quartet at the MLL1-NCP interface significantly reduced H3K4 methylation activity in an NCP-specific manner (10). However, it remains to be determined whether MLL1-NCP^mode 1^ represents an active conformation. Mutations of residues at key interaction interfaces in MLL1-NCP^mode 1^ were not functionally tested in a histone methyltransferase (HMT) assay (11). To fill this gap, we used our robust *in vitro* HMT assay to assess how unique interactions in MLL1-NCP^mode 1^ affect MLL1 catalysis on the NCP and recombinant H3. To this end, we first tested the function of the highly conserved _248_LVNR_251_ amphipathic loop in RbBP5 by introducing alanine mutations to L248/V249 or LVNR (LVNR→4A). As shown in Figure 2D and S6A, these mutations did not affect MLL1 activity on either the NCP or recombinant H3, suggesting that they do not contribute to MLL1 activity *in vitro*. Next, we deleted or introduced alanine mutations to key residues of the helical patch (3806-14) of the MLL1^SET^ domain (Figure 2C). As shown in Figure 2E, deletion of 3806-14 of MLL1^SET^ reduced H3K4 methylation on the NCP. Similarly, mutating M3812/L3814 to alanine also significantly reduced H3K4 methylation on the NCP (lane 3), which was partially ameliorated by the triple alanine mutant (3812/14/18A) for unknown reasons. However, these MLL1^SET^ mutants showed similar reduction of methyltransferase activity on recombinant H3 (Figure S6B), suggesting that they probably affect intrinsic activity of MLL1^SET^, instead of disrupting MLL1-NCP interactions. Finally, we found that mutating the arginine anchor, R3821, or the H2A acidic patch (NCP^APM^) also did not significantly affect MLL1 activity on the NCP. Taken together, our results show that the residues that are specific for MLL1-NCP^mode 1^ interactions do not contribute to NCP-specific activity of the MLL1 complex *in vitro*.

## DISCUSSION

Here we show two distinct binding modes of the MLL1-NCP^K4M^ complex *in vitro*. Together with our previous study (10), we find that only MLL1-NCP^mode 2^ is likely to be functionally active. In this mode, MLL1^SET^ resides above the nucleosome dyad, granting near symmetrical access to both H3 tails for optimal processivity (10). In contrast, the MLL1-NCP^mode 1^, where MLL1^SET^ binds across the nucleosome disc, is likely inactive since disruption of specific interactions between MLL1 and the NCP does not significantly alter MLL1 activity in an NCP-specific manner. Notably, MLL1-NCP^mode 1^ is similar to ySET1-NCP or ySET1-NCP^H2BK120ub^ complexes (12,13). The ySET1 complex binds across the nucleosome disc with the highly conserved _271_IINR_274_ loop of Swd1 anchoring on DNA near SHL 2.5. Mutation of the _271_IINR_274_ loop in Swd1 leads to complete loss of higher methylation of H3K4 in yeast (12) while its conserved counterpart, i.e. the _248_LVNR_251_ amphipathic loop in RbBP5, is dispensable for MLL1 activity on the NCP (Figure 2). Furthermore, the binding orientation of the ySET1 complex on the NCP enables extensive contacts between an unique arginine rich motif (ARM) in ySET1 and the acidic patch on the NCP (12,13). In the absence of H2BK120ub, ARM autoinhibits the SET domain, rendering it inactive on the NCP (12,13). The inactive state of ySET1 can be relieved by either H2BK123ub or removal of Spp1 (12,13). Notably, both ARM and Spp1 are not conserved in the MLL1 complex. Thus, despite similar binding orientations for the MLL1-NCP^mode 1^ and ySET1-NCP complexes, MLL1-NCP^mode 1^is probably subject to different regulations.

Our study highlights significant rotational dynamics of the MLL1 complex on the NCP. The MLL1 complex can rotate clockwise or counterclockwise with ASH2L (this study) and RbBP5 (Lee et al., paper in press) as anchors, respectively. As a result, the MLL1^SET^ deviates from the nucleosome dyad, profoundly impacting trimethylation activity on the NCP (10). Our study raises the question of why the MLL1 complex has higher rotational dynamics on the NCP and what are the functional implications. We envision several possibilities that await future studies. First, the observed MLL1 dynamics on the NCP may be due to the use of a minimal MLL1 core complex in the *in vitro* structural study. It is possible that additional MLL1 interacting proteins are needed to further stabilize the MLL1 complex in a specific conformation. We recently show that DPY30 is able to stabilize MLL1 binding in the active mode (MLL1-NCP^mode 2^) thereby significantly enhancing MLL1 trimethylation activity at gene promoters (Lee et al., paper in press). It remains to be determined whether other MLL1 interacting proteins are necessary to reduce MLL1 rotational dynamics *in vitro*. Alternatively, it is possible that linker DNA or adjacent nucleosomes are needed to further stabilize the MLL1 complex *in vitro*. Given that both the PRC2 and Rpd3S complexes require linker DNA for optimal chromatin binding and catalysis (19-21), it is possible that the nucleosome template assembled with linker DNA or an oligo-nucleosome array may reduce rotational dynamics of the MLL1 complex *in vitro*. Interestingly, the winged-helix motif in ASH2L is able to interact with DNA in a sequence-independent manner (22,23), which potentially allows for additional interactions with linker DNA. It would be interesting to investigate the structure of the MLL1 complex on di-nucleosomes or oligo-nucleosomes in the future. Finally, increased rotational dynamics of the MLL1 complex on the NCP is likely a unique feature for the MLL family enzymes in higher eukaryotes. It has been shown that MLL1 is able to mono- and di- methylate H3K4 at distal enhancers, but is also responsible for ∼5% H3K4me3 at gene promoters in cells (7,24). Studies in the past decade have underscored distinct distributions and functions of different H3K4 methylation states in higher eukaryotes (5,9,25-30). We speculate that high rotational dynamics of the MLL1 complex on the NCP may reflect a functional or regulatory requirement, enabling loci-specific regulation of the H3K4 methylation states. In this case, protein factors (e.g. DPY30) may regulate MLL1 activity by shifting the MLL1 binding modes and cementing MLL1 in a specific configuration. It would be interesting to examine whether proteins in the transcription machinery or in chromatin remodeling complexes are able to regulate MLL1 activity in this manner. In contrast to MLL family enzymes in higher eukaryotes, ySET1 complex does not display any rotational dynamics *in vitro* (12,13), consistent with the fact that ySET1 is more strictly regulated by H2BK123ub (31) and yeast generally lacks distal enhancers for transcriptional regulation (32). Interestingly, removal of Spp1 in the ySET1 complex, which provides additional anchor of the ySET1 complex on the NCP, derepresses SET1 activity on the NCP (12). Whether removal of Spp1 increases rotational dynamics of the ySET1 complex remains to be investigated in the future.

## EXPERIMENTAL PROCEDURES

### Protein expression and purification

MLL1 complex subunits (MLL1^SET, 3762-3969^, ASH2L^1-534^, RbBP5^1-538^, WDR5^23-334^, and DPY30^1-99^) and mutants were expressed using the pET-28a expression vector with N-terminal SUMO- and His_6_-tags (6). Deletion and point mutation plasmids were constructed using overlapping PCR and confirmed by sequencing. All proteins were expressed in BL21(DE3) *E. coli* strain in LB media. Cells were grown to OD_600_ at 37 °C until 0.6-0.8, and expression was induced by adding 0.4 mM IPTG. After 16-18 hours at 20 °C, cells were lysed by sonication and soluble lysate was collected by centrifugation at 32,000 x g for 30 minutes at 4 °C. After filtration through a 0.45 µm syringe, the soluble fraction was loaded and purified through a Ni-NTA metal-affinity column (Goldbio). After washes with 20 mM Tris, pH 8.0, 300-500 mM NaCl, 2 mM ®-mercaptoethanol, 10% v/v glycerol, and 10 mM imidazole (wash buffer), proteins were eluted with a stepwise imidazole gradient at 30, 60, 90, 120, 150, 210, and 300 mM. Fractions containing protein of interest were pooled and dialyzed overnight at 4 °C in presence of SUMO-tagged ULP1. Negative Ni-NTA purification was repeated to remove SUMO- and His_6_-tag, ULP1 and other bacterial impurities by collecting protein in flowthrough. Proteins were further purified on a HiLoad 16/600 Superdex 75pg or 200pg gel-filtration columns (GE Healthcare). To obtain stoichiometric MLL1^RWSAD^ complex, this complex was purified by combining equimolar amounts of MLL1^SET^, ASH2L, RbBP5, WDR5, and excess DPY30 and purified by HiLoad 16/600 Superdex 200pg gel-filtration column.

For histone purifications, full-length *Xenopus laevis* histones H2A, H2B, H3, H3K4M, and H4 were expressed and purified according to one-pot protocol (33). Briefly, histone constructs were transformed into BL21 (DE3), except for H4, which used C41 (DE3). After growing at 37 °C to OD_600_ of 0.6-0.8, protein expression was induced with 0.4 mM IPTG for 3 h (H2A, H2B, H3, and H3K4M) or 2 h (H4). Equimolar amounts of histones were combined and isolated from inclusion bodies and subject to octamer refolding (33). Octamer concentration was determined using UV at 280 nm. Reconstitution of nucleosome was conducted by combining 147 bp Widom 601 DNA and octamer in 1:1 molar ratio in high salt buffer using standard linear salt gradient method (34) overnight at 4 °C. Low salt buffer (20 mM Tris-HCl, pH 7.5, 1 mM EDTA, 1 mM DTT) was added via a peristaltic pump at ∼ 1 ml/min. Nucleosomes were then further dialyzed into long-term storage buffer (20 mM cacodylic acid, pH 6.0, 1 mM EDTA) overnight at 4 °C.

### *In vitro* histone methyltransferase assay

For *in vitro* HMT assay, 0.3 µM MLL1 complex was mixed with *S*-adenosyl-L-methionine and either NCP (1 µM) or recombinant H3 (0.1 µM) in 20 µL of HMT buffer (20 mM Tris-HCl, pH 8.0, 50 mM NaCl, 5 mM MgCl_2_, 1 mM DTT and 10% v/v glycerol) as previously described (35). The reactions were incubated at room temperature for 1hr and quenched by adding 20 µL 2x SDS-PAGE loading buffer.

### Western blotting

The histones were separated on a 15% polyacrylamide gel and transferred onto polyvinylidene difluoride membrane (PVDF, Millipore). The membrane was blocked in blocking solution, consisting of 5% milk in TBS buffer with 0.1% Tween-20 (TBST) and then incubated for 2 hours at room temperature with primary antibody. After washing 3 times with TBST, the membrane was incubated with the HRP-conjugated anti-rabbit secondary antibodies at room temperature for 1 h and developed using Pierce™ ECL Western Blotting Substrate (Thermo Fisher Scientific, #32106). The images were captured on ChemiDoc™ Touch Imaging System (Bio-Rad). The primary and secondary antibodies used in this study include: rabbit anti-H3K4me1 (Abcam, cat ab8895, 1:20000), rabbit anti-H3K4me2 (Millipore, cat # 07-030, 1:20000), rabbit-anti H3K4me3 (Millipore, cat # 07-473, 1:10000), and anti-Rabbit IgG Horseradish Peroxidase (HRP)-linked whole antibody (GE Healthcare, #NA934, 1:10000).

### Electrophoretic mobility shift assay

EMSA was conducted by incubating 0.1 µM NCP and with increasing concentration of the MLL1^RWSAD^ complex in 10 µL buffer. The protein/NCP mixture was then loaded onto a 6% 0.5x TBE gel and run for 1.5 hours at 150 V on ice. After the run, the gel was stained with 1:20,000 diluted ethidium bromide for 10 minutes and visualized by UV transillumination on Bio-Rad ChemiDoc Imaging System. Quantification was done using ImageJ software (36).

### CryoEM sample preparation and data collection

The MLL1-NCP samples were prepared using the GraFix method (37) as previously described (10). In brief, 30 µM of MLL1 complex was incubated with 10 µM NCP and 0.5 mM *S*-adenosyl-L-homocysteine for 30 min at 4 °C in the GraFix buffer (50 mM HEPES, pH7.5, 50 mM NaCl, 1 mM MgCl_2_, and 1 mM TCEP). The sample was centrifuged at 48,000 rpm at 4 °C for 3 h through a gradient 0-60% glycerol and 0-0.2% glutaraldehyde. After centrifugation, the crosslinked sample was quenched with 1 M Tris-HCl, pH 7.5 and glycerol was removed by buffer exchange. The sample at 1 mg/ml was applied onto Quantifoil R1.2/1.3 grids (Electron Microscopy Sciences) and the grid was plunge-frozen in liquid ethane using a Vitrobot Mark IV (Thermo Fisher Scientific) at 4 °C, 100% humidity and 4 sec blotting time. The cryoEM data was collected on the 300 keV FEI Titan Krios (Thermo Fisher Scientific) equipped with the Gatan K2 summit direct electron detector at a magnification of 130,000x in a counted mode. Each micrograph was imaged at the pixel size of 1.06 Å/pixel with the defocus range of -1.0 to -2.5 µm. A dose rate of 1.34 electrons/Å/frame with a total 40 frames per 8 sec was applied, resulting in an accumulated dose of 53.4 elections per Å^2^. A total of 2,377 movies were collected.

### CryoEM data processing and model refinement

Micrographic movie stacks were subject to MotionCorr2 (38) for whole-frame and local drift correction, and Contrast transfer function (CTF) was performed using CTFFIND4.1 (39). Micrographs with lower than 4.5 Å of the estimated resolution were excluded, which resulted in 2,377 micrographs. Particle picking was performed using Warp (40), and Warp picked a total of 808,836 particles The particles were extracted in RELION (41) and imported into cryoSPARC (42) for 2D classification. A total of 768,708 particles, after excluding bad classes from the 2D classification, were subjected to the first round of *ab initio* 3D classification into three classes (Supplemental Fig. 4). One of the three classes showed a clear density for the MLL complex and the NCP. This class of particles was then used for the second round of *ab initio* 3D classification into three subclasses. Two of the three classes seemed to maintain a well-defined cryoEM map of the MLL1 and NCP complex, but the binding patterns of MLL1 toward the NCP were distinguishable. Therefore, two subclasses were subjected for subsequent heterogeneous refinement independently. Two particle sets were further exported to RELION for additional 3D classification. For each class, the focused 3D classification was performed at the MLL1^RWSAD^ region without alignment (35 cycles, T=4, binary mask: 10 pixels/soft mask: 10 pixels). The best behaving class selected from Class 01 (30,847 particles, Mode 2) and Class 03 (30,322 particles, Mode 1) were subjected to 3D auto refinement. After CTF-refinement and particle polishing, each class was further refined and post-processed to a resolution of 4.03 and 4.76 Å, respectively. The resolution of all structures was estimated by RELION with Fourier shell correlation (FSC) at the criteria of 0.143. To build the atomic model of the MLL1-NCP^K4M, mode 1^ and MLL1-NCP^K4M, mode 2^, the structures of MLL1-NCP (PDB ID: 6KIX and 6PWV) were, respectively, used for rigid-body fitting. The real-space refinement using PHENIX (43) was performed after the rigid-body fitting. For the model validation, MolProbity (44) was used, and the map and model FSC curves were calculated using phenix.mtriage in the PHENIX program package (Figure S5C and D). Statistics of data collection, refinement, and validation are summarized in Table 1.

## DATA AVAILABILITY

The accession numbers for the MLL1- NCP^K4M, mode 1^ and MLL1- NCP^K4M, mode 2^ structures are PDB: 7MBM [https://www.rcsb.org/structure/7MBM] and EMDB: EMD-23738 and PDB: 7MBN [https://www.rcsb.org/structure/7MBN] and EMDB: EMD-23739, respectively.

## ACKNOWLEDGEMENT AND FUNDING

This work is support by the NIGMS grant (GM082856) to Y.D and the NCI grant (CA250329) to Y.D and U.S.C and National Cancer Institute Cancer Center Shared Grant award P30CA014089 for Norris Comprehensive Cancer Center at University of Southern California. A.A. is supported in part by the MERIT fellowship from the Rackham graduate school at University of Michigan. This research was, in part, supported by the National Cancer Institute’s National Cryo-EM Facility at the Frederick National Laboratory for Cancer Research under contract HSSN261200800001E.

## AUTHOR CONTRIBUTIONS

A.A. designed and performed biochemical experiments and wrote the manuscript. Y.T.L performed experiments with acidic patch mutants. S.H.P. performed the cryo-EM studies under the supervision of U.-S.C. Y.D. and U.S.C. supervised the overall study and wrote the manuscript. All authors read and approved the manuscript.

## CONTACT FOR REAGENT AND RESOURCE SHARING

Further information and requests for resources and reagents should be directed to and will be fulfilled by the Lead Contact, Yali Dou.

